# CRISPR/Cas9-mediated mutagenesis of the Asian Citrus Psyllid, *Diaphorina citri*

**DOI:** 10.1101/2023.05.05.539615

**Authors:** Duverney Chaverra-Rodriguez, Michelle Bui, Cody L. Gilleland, Jason L. Rasgon, Omar S. Akbari

## Abstract

The most devastating disease affecting the global citrus industry is Huanglongbing (HLB), caused by the pathogen *Candidatus Liberibacter asiaticus*. HLB is primarily spread by the insect vector *Diaphorina citri* (Asian Citrus Psyllid). To counteract the rapid spread of HLB by *D. citri*, traditional vector control strategies such as insecticide sprays, the release of natural predators, and mass introductions of natural parasitoids are used. However, these methods alone have not managed to contain the spread of disease. To further expand the available tools for *D. citri* control via generating specific modifications of the *D. citri* genome, we have developed protocols for CRISPR/Cas9-based genetic modification. Until now, genome editing in *D. citri* has been challenging due to the general fragility and size of *D.citri* eggs. Here we present optimized methods for collecting and preparing eggs to introduce the Cas9 ribonucleoprotein (RNP) into early embryos and alternative methods (ReMOT Control) for injecting RNP into the hemocoel of adult females for ovarian transduction. Using these methods, we have generated visible somatic mutations, indicating their suitability for gene editing in *D. citri*. These methods represent the first steps towards advancing *D. citri* research in preparation for future genetic-based systems for controlling HLB.

## Introduction

The Asian citrus psyllid (ACP), *Diaphorina citri* (Hemiptera: Liviidae), is the vector of the bacteria *Candidatus Liberibacter asiaticus* (CLas), the etiological agent responsible for Huanglongbing (HLB) disease in citrus (1)). HLB is the most destructive citrus disease worldwide, causing fruit from infected trees to become small, misshapen, and discolored with an unpleasant bitter and acidic flavor, making them unmarketable (2). The damage caused by HLB is extensive, decreasing fruit production by 57% in Florida alone from 2004–2019 and causing an estimated $1.7 billion in production losses between the harvest seasons of 2006–2007 and 2010–2011 (3). The initial symptoms of the disease do not appear for months and at first resemble nutrient deficiencies, resulting in HLB infections remaining undiagnosed for long periods and providing stable reservoirs for the pathogen (4). As an example of its rapid spread, HLB reached the Western Hemisphere by 2004 (5,6), and by 2013, every grove in Florida was considered infected(3). Currently, there is no commercially available cure for trees infected with HLB (7,8).

HLB is mainly transmitted by the ACP vector, but it can also be spread from tree to tree through grafting diseased branches (9). Thus, the main method for reducing the spread of HLB among commercial groves is a three-pronged system that includes (i) removing infected trees serving as HLB reservoirs, (ii) planting certified healthy plants, and (iii) applying insecticides to control *D. citri* vector populations (10). Although this method has seen moderate success, it doesn’t completely stop CLas, so HLB continues to spread. In response, numerous methods for *D. citri* population control have been proposed and field tested. These methods range from new insecticides to the release of natural predators and parasitoid wasps. These methods still have their issues including the development of insecticide resistance, the lack of efficacy and penetration of control, and the expensive costs needed for continued application (11–15).

To further expand the available methods of *D. citri* control, we looked into genetic-based control methods similar to those used in other insect vectors of disease and agricultural pests, such as *Aedes aegypti* and *Drosophila suzukii*. Gene editing tools based on CRISPR/Cas9, such as gene drives, pathogen-resistant effectors, and precise guided Sterile Insect Technique (pgSIT) (16–18) have the possibility to significantly contribute to the management of HLB. However, implementing these tools in *D. citri* has proven challenging mainly due to the complex methods required for delivering Cas9 or DNA into early embryos. *D. citri* lay small eggs, 0.3 mm long x 0.14 mm diameter, attached to plant tissues with a basal stalk or pedicel, which facilitates essential water exchange with the plant. Clusters of eggs can be laid at the center of, or embedded within the tightly guarded plant flush (1,19). These features make the eggs extremely difficult to reach for collection without affecting their survival. In addition, the delicate nature of the pre-blastoderm embryo further complicates mass collection, egg alignment, injection and transfer, leading to a high fatality rate by manipulation alone. Thus, previous attempts at genetic manipulation of *D. citri* using traditional embryonic microinjections have shown to be difficult and unsuccessful (20). Therefore, innovative strategies for embryonic microinjection that reduce egg manipulation or completely avoid egg removal from the plants may be critical for the successful delivery of CRISPR/Cas9 in *D. citri*.

Alternatives that bypass embryonic microinjection for delivering CRISPR/Cas9 for germline mutagenesis have been developed and tested in different arthropod species (21–26), though they have yet to be adapted and optimized for *D. citri*. Briefly, Receptor-Mediated Ovary Transduction of Cargo (ReMOT Control) delivers CRISPR/Cas9 ribonucleoprotein (RNP) into insect ovaries by fusing the RNP to a peptide ligand derived from yolk proteins. ReMOT Control has been used to create heritable gene edits in the germline of several mosquito species, the silverleaf whitefly, the jewel wasp, *Tribolium* beetles, brown marmorated stinkbug, and the blacklegged tick (21,23,24,27–29). Alternatively, Branched Amphiphilic Peptide Capsules (BAPC) were used recently to improve the delivery of CRISPR components into adult ovaries to facilitate heritable gene editing in *D. citri*. BAPC-assisted Cas9 RNP delivery caused targeted disruption of the Thioredoxin-2 gene (*trx-2*) in the germline of fifth-instar nymphs and adult female ACP (20). These results encourage the exploration of additional female injection techniques such as ReMOT Control in *D. citri*.

Here, we demonstrated for the first time successful CRISPR/Cas9 gene editing in *D. citri* using embryonic microinjection and ReMOT Control. To ensure gene modification, we targeted genes with visible mutant phenotypes, *white* (*w*) and *kynurenine hydroxylase (kh).* We identified the location of these genes in the *D. citri* genome to design targeting gRNAs, developed and tested Cas9-RNP delivery methods into embryos and females, and validated the phenotypic and genotypic changes upon gene disruption. The development of these protocols is crucial to apply the techniques necessary for transgenesis of *D. citri,* and to create genetic control systems for the reduction of HLB.

## Results

### Assessment of target sites and sgRNA design

To demonstrate effective genomic DNA targeting, we chose to target two genes conserved in several insect species known to produce visible phenotypic changes in eye color when disrupted, *w* and *kh* (*30–33*). Mutants for either gene typically display a reduction of pigmentation in their eyes. We identified the genomic sequences for *w* and *kh* using the latest version of the *D. citri* genome (*Diaphorina citri* version 3.0, Diaci v3.0; https://citrusgreening.org/tools/blast?db_id=17) by comparing it to the known protein sequences of the *D. melanogaster w* and *kh* genes using tBlastn (NCBI). *D. melanogaster w* aligned to genomic sequences found in 6 of the 13 *D. citri* chromosomal-length scaffolds (DC3.0sc01 to DC3.0sc13), and 15 out of 31 hits were located within scaffold DC3.0sc02 (**Table S1**). We downloaded this sequence (total of 47,354,213 bp) and annotated the aligned sequences using *D. citri* mRNA and protein databases. The ortholog contained 12 exons and aligned with the mRNA sequence ID XM_026820729.1 on NCBI (2828 bp), corresponding to an ABC transporter protein predicted to be the *w* gene, sequence ID XP_026676530.1 (**Fig. 1A**). The *D. citri kh* gene was located at scaffold DC3.0sc01 between nucleotides 9,672,699 and 9,676,291. The annotated ortholog contained 5 exons and aligned with the mRNA sequence ID XM_008469982.3 on NCBI (1227 bp), corresponding to a kynurenine 3-monooxygenase-like gene, protein sequence ID XP_008468204.1 (240 amino acids), that partially aligns to *D. melanogaster kh* protein sequence (465 amino acids). We blasted XM_008469982.3 against the *D. citri* proteins in NCBI and found an additional sequence, ID KAI5750130.1 (527 amino acids), that had higher coverage and aligned better with *D. melanogaster kh* protein (**Fig. 1A**). We used this protein to annotate the *kh* gene in the genomic sequence at scaffold DC3.0sc01. We then identified potential sgRNA target sites marked with a short protospacer adjacent motif (PAM; 5’-NGG-3’) across exons 1–6 of the *w* gene and on exons 7 and 8 of the *kh* gene. We designed primers across both genes and validated the predicted target regions in our *D. citri* colony (**Supplementary Information, Table S2**). We generated sgRNAs for target regions that did not show significant nucleotide variation and these were tested for *in vitro* cleavage using the PCR product of the target gene (**Fig. 1B**, **Figures S1–S3**). The final list of sgRNAs contained only those with high targeting efficiency and cleavage rates, resulting in three sgRNAs targeting sites on exons 2, 3, and 6 of the *w* gene and two sgRNAs targeting two sites on exon 7 of the *kh* gene (**Table S3**).

**Figure 1.**
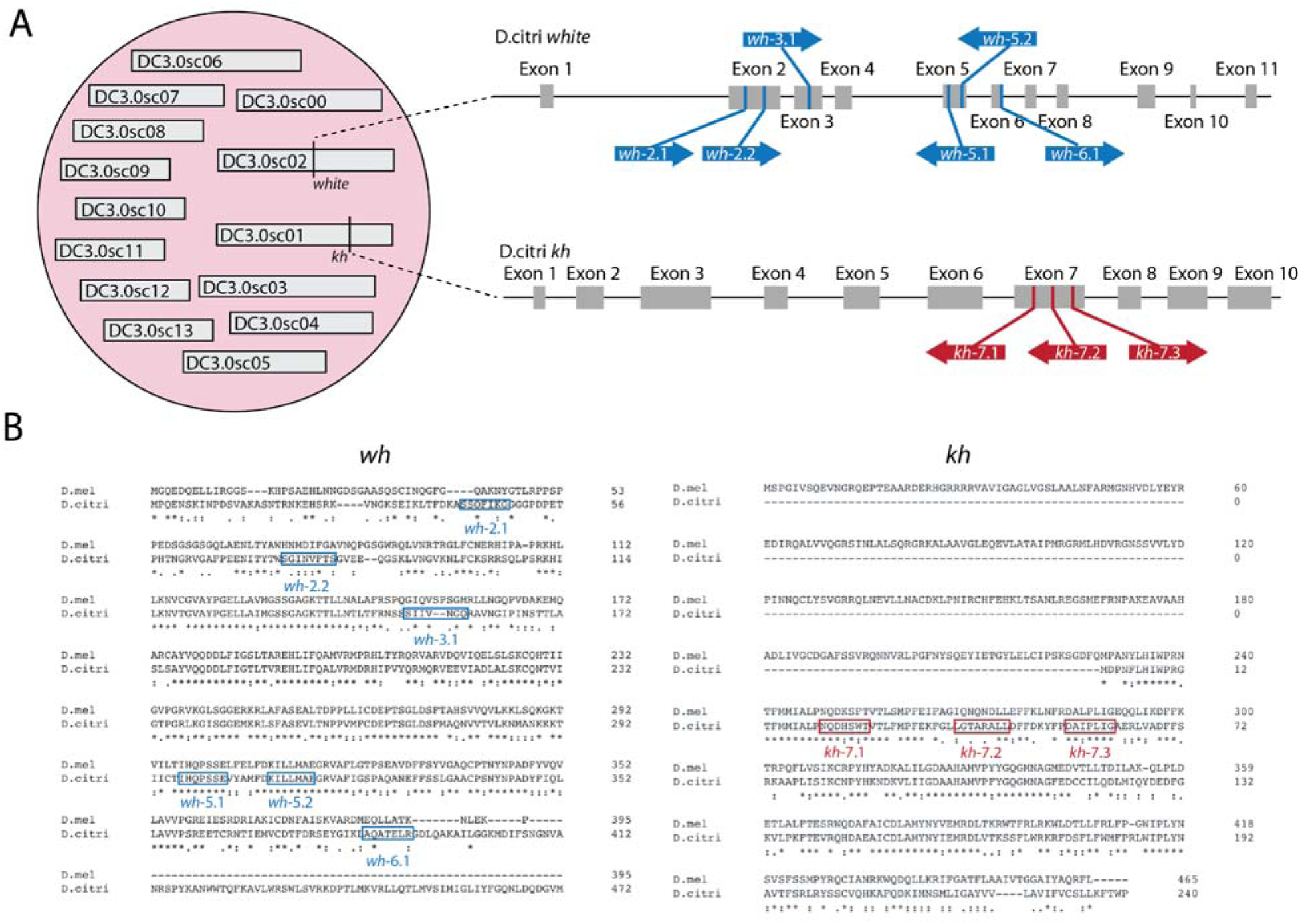
Molecular confirmation of *white* and *kh* genes in *D. citri* and selection of sgRNAs. A) The *w* and *kh* genes were located within the *D. citri* genome using tBLASTn with the *D. melanogaster* orthologs of both genes. *W* was within the DC3.0sc02 scaffold, and *kh* was in DC3.0sc01. Six sgRNAs were designed to target *white.* Three sgRNAs were designed to target *kh.* B) SgRNAs targeted regions in *D. citri* with conserved and non-conserved nucleotide and amino acid sequences compared to the DNA and protein sequences resulting from *D.melanogaster w* and *kh* genes, respectively.

### Development of protocols for *D. citri* embryonic microinjection

We found multiple bottlenecks to carry out efficient injections on *D. citri* embryos, these included hard-to-reach and remove eggs, needle clogging, and reduced embryo survival after injection. To address these challenges, we developed two approaches. Our first approach (Protocol-1) used the standard embryo injection technique in which eggs are collected, aligned on a microscope slide, injected, and incubated until eclosion (**Fig. 2A**) (34,35). We tailored our methods to accommodate *D. citri* biology aiming to reduce egg mortality by optimizing three critical features: a technique for gentle egg collection and alignment, methods to avoid needle clogging, and post-injection conditions for embryo incubation and nymph hatching (**Supplementary Protocol 1, Fig. S3**). In brief, we collected embryos from curry plants, *Murraya koeniigi,* using a dissecting teasing needle with a curved bend at the tip which allowed us to pluck the egg’s pedicel from the plant. Embryos were aligned and injected on filter paper soaked in sterile ultrapure water. To address needle clogging, we prepared 10 needles before each session, loading them with 3–4 ul of the mix when needed, and we adjusted the compensation pressure parameter (Pc) in the femtoJet(R) to maintain a consistent volume of solution injected. Once all eggs were injected, they were transferred to the hatching media reported by (36) to provide nutrients and maintain humidity and isolation from contaminants in the external environment. Eggs were transferred to fresh plates every two days to reduce mold growth until nymphs hatched, at this point these were moved to a new plant for feeding (**Fig. 2A**).

**Figure 2.**
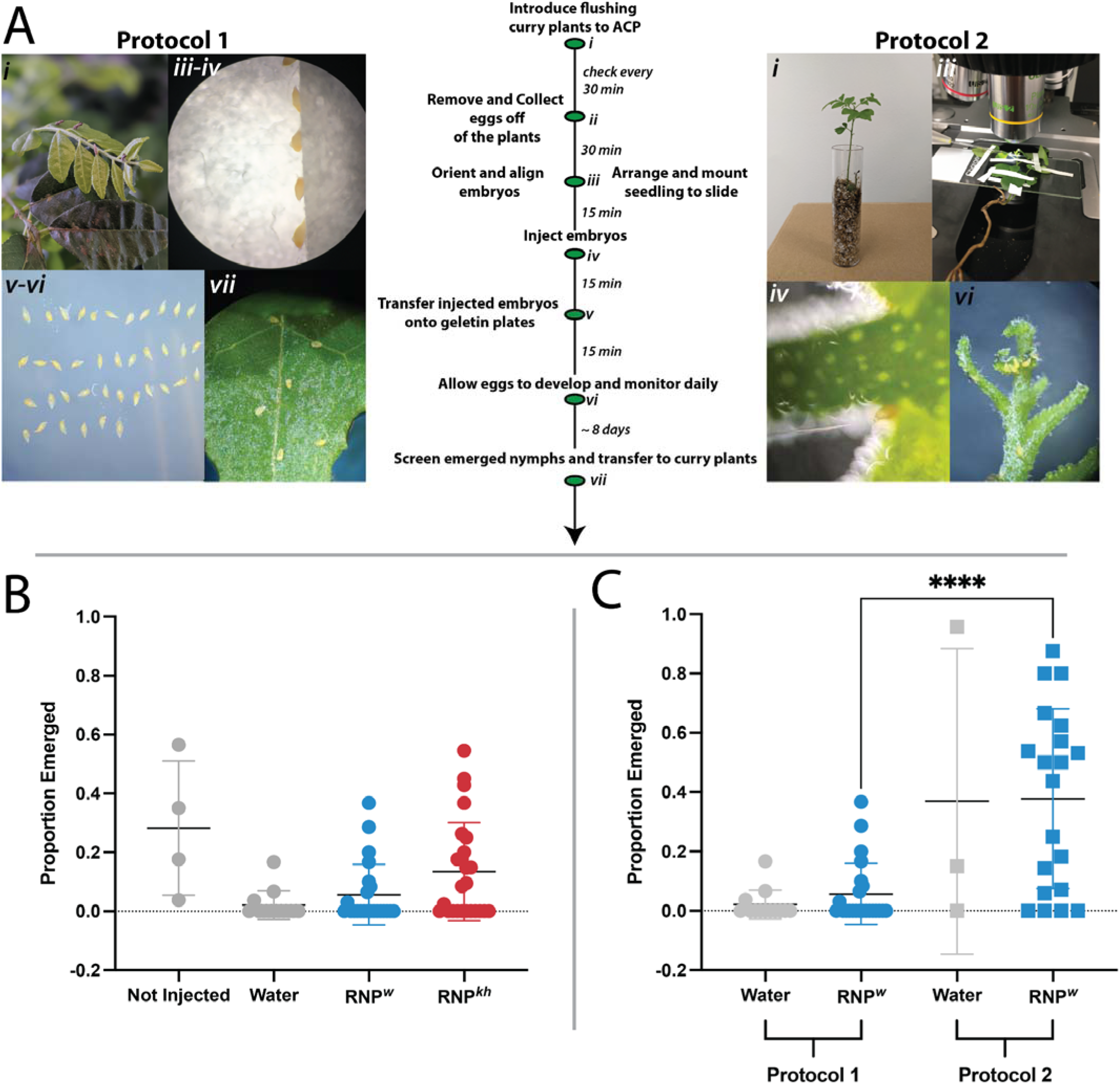
Injection protocols and observed rates for emerging nymphs from injected eggs. A. Schematic and timeline of embryonic injections using Protocol 1 and Protocol 2. B. Proportion of nymphs emerged under protocol 1. Eggs were removed from the plant and either not injected or injected with ultrapure water, RNP*^w^*, or RNP*^kh^*. C. Proportion of nymphs emerged comparing Protocol 1 (removed from plants) and Protocol 2 injected while remaining on the plant. (****p value <0.0001, unpaired t-test).

To determine the effects of Protocol-1 on hatching rates we evaluated several treatments: unmanipulated embryos that remained on the plants (N=108), embryos removed from the plant, aligned and incubated on hatching media (N=109), embryos aligned, injected with water (N=302), with RNP targeting the *w* gene (RNP*^w^,* N=633) or with RNP targeting the *kh* gene (RNP*^kh^*, N=949) incubated on hatching media. First, unmanipulated eggs that were left on plants as a control hatched at high rates (53.6%), though not all progressed to adults (25%) (**Table 1**). Removing and manipulating eggs reduced hatching to 33.9% compared to the hatching rates of embryos that were left on the plant (t-test = 1.2, df = 8, p-value =0.2646). Furthermore, removing eggs and injecting them with ultra-purified water reduced survival to 1.7% (t = 4.142, df = 15, p-value = 0.0009). Injecting RNP*^w^* and RNP*^kh^* resulted in 6.2% and 11.3% survival respectively (**Fig. 2B**). In addition, 1.4% of embryos and nymphs from injections with RNP*^w^*(sgRNA 2.1) showed altered phenotypes such as lack of pigmentation in one of the developing eyes or changes in body color that correlated with sequences consistent with mutations in the target site, this was not observed on eggs or nymphs hatching from RNP*^kh^* injections (**Table 1, Fig. S4**). None of the nymphs hatched from eggs injected with RNP*^w^* reached the adult stage, and thus, we were not able to set up crosses to demonstrate the heritability of these modifications (**Table 1**). Overall, these results confirm that delivery of RNP*^w^* into *D. citri* embryos using the developed protocols in this study produces phenotypes and genotypes consistent with mutations on the *w* gene but not on *kh*. However, lower survival rates affected our chances to demonstrate the heritability of the mutations. Therefore, aiming to improve survival rates for both nymph and adult stages, we developed a second protocol (Protocol-2) to inject *D. citri* embryos with RNP*^w^*while still attached to the flushes of curry seedlings (**Fig. S3, Supplementary Protocol 2**).

Protocol-2 involved using 14 *M. koeniigi* seedlings for *D.citri* females to lay eggs during 1-2 hours. At this stage, seedlings were uprooted, mounted on a glass slide to inject the eggs on the microscope, and planted back into the soil. We observed 100% seedling survival and scarce evidence of plant wilting after the entire process. This method required preliminary observations of the location of each egg, trimming the leaves that did not have eggs, and carefully taping the plant to the glass slide while keeping the eggs exposed in a proper angle for injection (**Fig. 3**). Successful injections (those in which injected solution entered the embryo creating a movement of the cytoplasm) typically occurred on eggs located at the axial branches, or on those located at the extremes of a row, given the physical support and stability these provide when the needle is pressed against the egg (<u>Supplementary Video 1</u>). Using this protocol, we independently injected groups of embryos with water (N=70) or RNP*^w^*(N=101) resulting in hatching rates of 32.9% and 33.6% respectively (**Table 1, Fig. 2C**). About 12% of G_0_ nymphs hatched from eggs injected with RNP*^w^* reached the adult stage, but none of them showed altered eye phenotypes or genotypes. After allowing these G_0_ adults to cross with each other we did not find any altered phenotype in their G_1_ progeny. Finally, a statistical comparison for RNP*^w^*injections showed significant differences on hatching rates between protocol 1 and protocol 2 (t = 4.781, df = 41, p-value <0.0001, unpaired t-test). These results showed that hatching rates and survival of injected embryos to the adult stage can be greatly improved by injecting eggs attached to the plants. Overall, our results confirm that delivery of RNP*^w^* into *D. citri* embryos using the developed protocols in this study produces phenotypes and genotypes consistent with mutations on the *w* gene.

**Figure 3.**
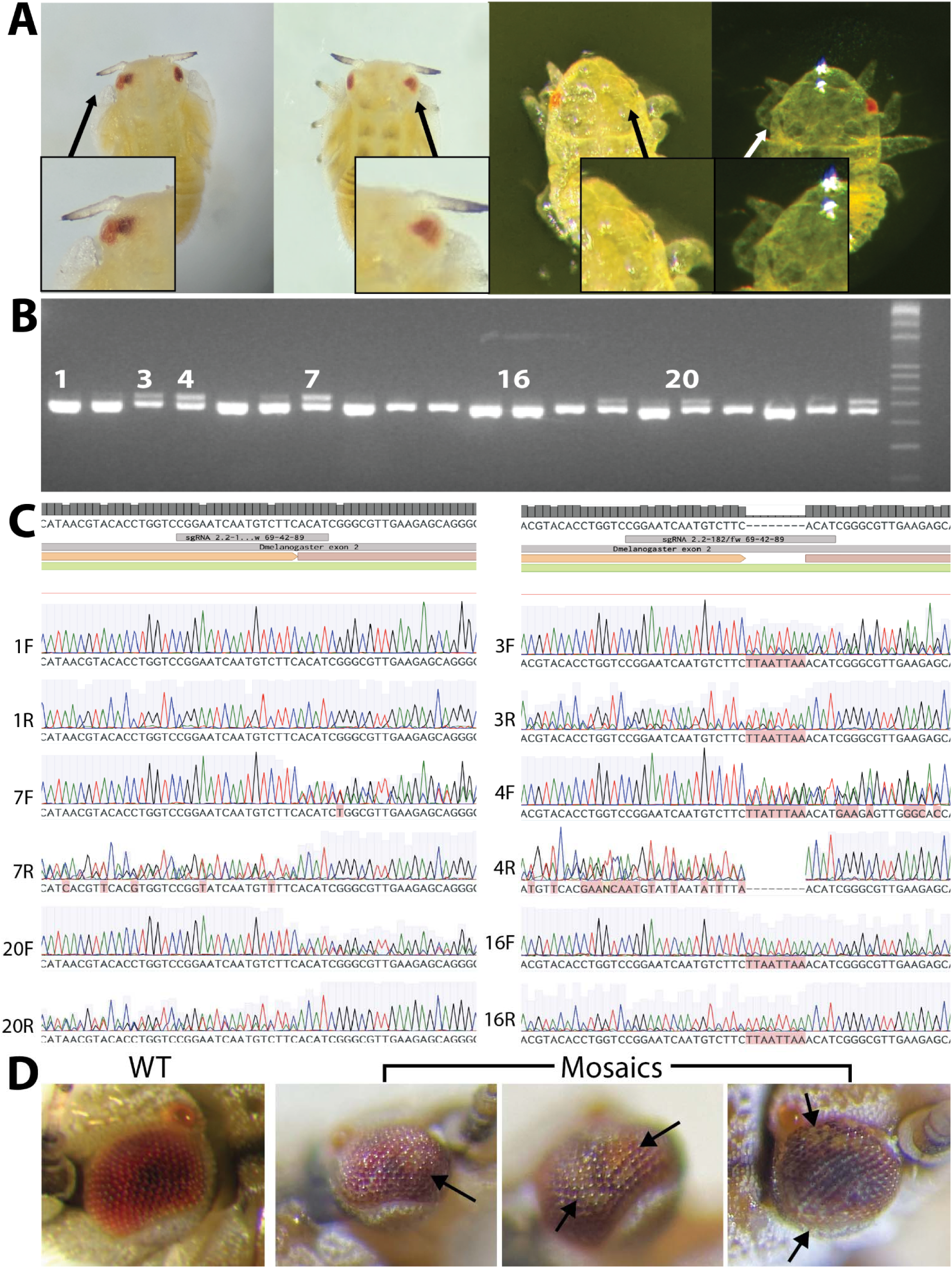
Generation and confirmation of *white* gene mutants. A) To search for potential mutant phenotypes, emerged nymphs were screened under a stereoscope. B) To visualize possible INDELs, PCR samples were run on a 4% agarose gel. C) Sanger sequencing confirmed INDELs within the target region. D) Adult mosaic KO phenotypes, arrows point to patches of white ommatidia.

### *D. citri* mutagenesis via adult female microinjections

To identify the optimal conditions to deliver active RNP*^w^*components into *D. citri* female ovaries, we tested ReMOT Control and BAPC. First, we injected five *D. citri* females with the fusion protein P2C-mCherry and observed intense red fluorescence in developing oocytes within 24h of injection (**Fig. S1C**) indicating that the ovaries were able to transduce the fused P2C protein. We injected groups of 7-10 days-old *D. citri* females and placed them in a cage containing plants for oviposition (**Fig. S3**). We screened the G_0_ progeny for changes in eye color and determined the rate of germline mutagenesis by screening the G_1_ or G_2_ offspring derived from outcrossed G_0_ mutants. Injecting P2C-RNP*^w^* at concentrations between 1000-2000 ng/uL protein and 800-2000 ng/uL sgRNA, resulted in >50% female survival regardless of the delivery method utilized (**Table 2**).

We observed a low rate of visible eye mutants with ReMOT Control (0.5-10% of the total G_0_ progeny from injected females) using sgRNAs 2.1, 3.1. and 6.1. Somatic mosaicism was detected in the eyes of two nymphs, two females and one male (**Fig. 3A, 3D**). The modified eyes of the G_0_ presented differences in ommatidia with loss of pigmentation from most of the eye or in patches distributed randomly (**Fig. 3B-C**, <u>Supplementary video 2</u>), consistent with results for somatic mosaicism in the *w* gene observed in other insect species (21). We collected individuals with mutant phenotypes, isolated their genomic DNA, and sequenced the target region. We observed indels within the target region of some individuals but for others, we observed overlapped secondary peaks and proceeded to carry out ICE analysis which showed that all the mosaic individuals had mutations on their target site (**Fig. 3B-C**). The two mosaic nymphs died before molting to the second instar, preventing us from identifying the heritability of the eye alteration. However, we independently crossed each of the mosaic G_0_ females to 10 wild-type males and the G_0_ mosaic male to 10 wild-type females and also intercrossed their G_1_ progeny. None of the G_1_ or G_2_ progeny from the mosaic male or females scored phenotypically or genotypically as a mutant. Interestingly, from ReMOT injections we also found two adult G_0_ males displaying non-typical light-orange/pink-colored bodies and gray eyes. These traits were not observed in any of the non-injected control groups or the laboratory colony (**Fig. S5**). Sequences from these individuals did not show any alteration on the sgRNA target site and we hypothesized whether these could be off-target mutants. We thus proceeded to cross each of these males independently to >10 females, producing eggs that did not develop into nymphs, which suggests the males were sterile. Overall, our data demonstrate that ReMOT Control can be used to produce somatic *white* gene mutations on *D. citri*.

On the other hand, two out of four of the BAPC experiments experienced massive death of G_0_ eggs and nymphs (~90%) producing few G_0_ adults which did not show mutant phenotypes. When we analyzed a sample of sequences from randomly picked dead G_0_, we found altered sequence peaks that expanded around the target sites (**Fig. 4**). Moreover, control injections consisting of direct injections of Cas9-RNP*^w^*without the transfection reagent (and without the peptide P2C) resulted in 12 out 136 G_0_ nymphs showing altered eye phenotypes (8.8% G_0_). Seven of these (5.15% of G_0_) showed sequences corresponding to *indels* in the target site for the sgRNA2.1 (**Table 2**). Altogether, commercially available Cas9 RNP delivered through BAPC produced alterations on the *white* gene with low penetrance. However, removing the BAPC, the P2C ligand, or the endosomal escape reagents from the RNP mix increased the number of mutants observed per injected female.

**Figure 4.** 2 nymphs, 3 mosaic adults from ReMOT - has CRB with 3 adults.

## Discussion

Here, we demonstrate targeted somatic mutations in *D. citri* using CRISPR-based technologies delivered via embryonic and adult female microinjections. Achieving site-directed mutagenesis in *D. citri* using traditional embryonic microinjection methods has been elusive. To accommodate *D. citri* features for genetic manipulation using Cas9-RNP, we have developed two protocols to collect and inject embryos with or without removing the eggs from their plant substrates.

Protocol-1 was based on embryonic injection techniques validated in model organisms such as *Ae. aegypti,* but also in non-model hemipterans such as *Nilaparvata lugens*, *Lygus hesperus, and Pyrrhocoris apterus* (33,37,38). Because this protocol has the eggs removed from the plant, it has multiple advantages such as facilitating that every egg aligned on the glass slide can be injected. In addition, the host plants can be immediately reused to collect additional rounds of eggs within the same injection session. However, the damage caused by both removing the eggs from the flush and the subsequent manipulation needed for alignment and incubation, may lead to lower rates of survival and thus reduces the chance of generating a mutant. Thus, we proceeded to systematically identify the bottlenecks of our protocol, and improved the techniques regarding egg collection, avoidance of needle clogging, and treatment of eggs post-injection. First, ACP females start laying eggs once mated around 2-3 days of emerging (39), thus egg collection required synchronizing the laboratory colony and curry plant rearing to produce a considerable number of gravid females and flushing plants amenable for oviposition on each injection session, we considered the number and age of females and males for the oviposition cages, the timing of egg collection and the size and number of flushes on each plant. Second, the freshly laid eggs are fragile and sensitive to the pressure needed to pluck the eggs from the plant substrate without damaging the pedicel (36). We had to be gentle at egg removal and alignment, making this process very challenging and time-consuming. Thus, we included an intermediate step in which eggs were transferred from the plants to agarose plates, agarose is soft and pliable, allowing the embryo pick to pass through instead of needing to rotate and potentially break the egg. We then used a fine paintbrush to gently align the eggs on the glass slide. Third, during injection, the dense cytoplasm of *D. citri* embryos clogged our needles constantly. The cytoplasm would cover the tip of the needle and if left to dry it would harden and create a clog. We found no way to avoid this issue totally, however, it can be minimized by adjusting the values of Pc on the Femtojet to levels in which small droplets of solution are constantly pushed out the needle, countering the embryo pressure.

Using our optimized Protocol-1, we demonstrated for the first time CRISPR/Cas9-induced phenotypes and genotypes after injecting RNP^w^/sgRNA 2.1 into *D. citri* embryos. However, we were not able to demonstrate the heritability of these mutated genotypes because the nymphs that hatched on gelatin media died a few days after being transferred to host plants. Santos-Ortega & Killiny (2020) observed that nymphs hatched on this media never molted to the second instar and that transferring nymphs to *Citrus macrophylla* plants allowed 56% of nymphs (n=5) to survive to adulthood. The authors suggest that nutrients and conditions on the plant may be required for nymphs to molt to the next instar. It may be a plausible explanation that the microenvironment created by curry plants impacts the adjustment of *D. citri* nymphs differently. Also, it is likely that the reduced ability of nymphs to adjust to curry plants might be related to fitness costs derived from the Cas9-induced *w* mutations. It was demonstrated recently in the hemipteran *O. fasciatus* that homozygous mutations on the *w* gene are lethal (30).

Our experiments showed that manipulation of embryos resulted in 33% of hatching while injecting embryos with RNP reduced hatching between 6% to 11%; these rates are comparable to those reported in recently developed protocols for non-model arthropods like ticks (4-12%) or hemipteran species such as *N. lugens* (2,9-3.5%) *or L. hesperus* (8.8-27.5%) (33,37,38,40). We believe that survival rates may be increased with further optimization of factors related to egg manipulation that affect mortality such as embryo collection or post-injection treatment of eggs. However, there is evidence that the connection of the egg pedicel to the plant is necessary during *D. citri* embryo development, and this may impose constraints on embryo survival rates for injection methods that require egg removal (Burckhardt 1994). Therefore, we worked on finding alternative methods for injecting the eggs while still attached to the plants using RNP*^w^*.

Protocol-2 provides the first successful approach reported for injecting *D. citri* eggs attached to plants, from which new improvements can be built by users. The greatest limitation of this method is that the final number of injected eggs per plant is very low, which makes it inefficient compared to the amount of time and effort required for the removal of eggs from locations that are difficult to impossible to reach with the needle. Furthermore, since the eggs had to remain on the plants, this method only allowed for a single use of the plant until the injected eggs hatched, requiring a high number of available seedlings for egg collection in a session. In our case, we used the HLB-resistant *M. koeniigi* as host plants for *D. citri,* we were able to obtain seeds only once a year at the beginning of the fall, attaining the first cohort of seedlings suitable for egg laying in December-January and the last one at the end of February. This short time period constrained our chances to test various treatments and optimize our protocols. Ideally, this may be improved by keeping a large number of seedlings with amenable flush shoots for oviposition during the whole year. Growing seedlings of other host plants with accessible seeds for the whole year could be a strategy. Alternatively, the production of plant shoots from vegetative or somatic *in vitro* propagation from these host plant species could also be explored (41–44). On the other hand, although Protocol-2 resulted in fewer injected eggs compared to Protocol-1, it drastically increased the hatching rates of eggs injected with RNP*^w^* (6% vs 34%), as well as the percentage of hatched G_0_ nymphs that reached the adult stage (0 vs 28%). However, none of these G_0_ adults or their G_1_ progeny showed the phenotypes or genotypes associated with *w* mutations observed in Protocol-1. This may be partially explained by the quality of the injections on the plant, which were more difficult to complete compared to injections of the eggs removed and aligned on a slide; as well as by a reduced efficiency of mutagenesis associated with fewer eggs injected. Due to the complex ways eggs are laid on the plants and the lack of rotational flexibility of the microscope stage used to move the plant piece, injections on the plants were difficult and time-consuming, features that may discourage its use by researchers. However, our results confirmed that embryo survival was greatly improved by injecting eggs located on plants, supporting our hypothesis that egg removal from the plant and their manipulation increases mortality. Further optimization of this method may facilitate the injection of plasmids for expression or transgenesis of *D. citri*.

In addition to embryonic injections, delivery of Cas9 protein to oocytes has also been achieved through injections into the adult female hemocoel to encourage transduction into the ovaries (1,3,4). We tested two different strategies: ReMOT Control and BAPC. For both methods, injections were performed at the first two sternites in the abdomen with an RNP mix containing concentrations around 2000ng/uL of P2C-Cas9, 1000 ng/uL of sgRNA and 1mM saponin for ReMOT Control, and 1000 ng/uL Cas9, 800 ng/uL sgRNA and 1200 ng/uL of BAPC. Females tolerated the injections well, however, the number of G_0_ observed after injection varied drastically compared to those of the controls injected with Cas9/sgRNA only or those non-injected. It is likely that components such as saponin or BAPC may have an impact on *D. citri* G_0_ development. For all our experiments we used saponin as an endosomal escape reagent (EER), at a concentration of 1mM. EERs facilitate the release of the RNP from endosomes once it is transduced into the oocytes (45). Saponin has been used successfully as an EER on *Ae. aegypti, An. stephensi,* and *Culex pipiens pallens* (*24,27,28,46*). However, in *N. vitripennis,* increased concentrations of saponin had impacts on the number of females that lay eggs and the number of larvae that entered diapause (28). In addition, saponin was inhibitory for editing during ReMOT Control in the hemipteran psyllid *Bemicia tabaci* (23). On the other hand, the G_0_ nymphs from the four experiments with BAPC suffered from drastic mortality in the first week after hatching without apparent reason. Compared to injections reported in *N. vitripennis,* our BAPC injection mix contained 2-3 times the concentration of each component because *D. citri* females are bigger than *N. vitripennis.* However, it may be possible that these concentrations may become detrimental to the nymph’s survival. Also, female and nymph mortality could be the result of plant quality, temperature, and humidity conditions rather than injection media. To avoid this, plants should be observed daily making sure that nymphs have the resources necessary for development. We did not test the effects of EER or different concentrations of BAPC in the fecundity of injected females or the survival of G_0_ nymphs, future work may benefit from carrying out these experiments.

We observed visible somatic mosaicism for the *white* gene of *D. citri* using ReMOT Control. The recovered adult mosaic mutants did not inherit the altered genotypes to their G_0_ or G_1_ progeny because these mutations were not in the germline. Nevertheless, the gene editing efficiency (percentage of mutant G_0_ out of the total hatched G_0_ = 1.2%) is comparable to that obtained for the jewel wasp *N. vitripennis* (0.75%), the ticks *Ixodes scapularis* (4.1%) and mosquitoes such as *Culex quinquefasciatus* (0.4%), *Ae. aegypti* (1.5%) and *An. stephensi* (3.7%). For some of these species, ReMOT Control has been proven useful to produce heritable genetic modifications, and thus, it may be required to optimize certain components of the injection protocol to obtain and isolate the germline mutants (24,27,28,46). We noticed that the rates of gene editing and mosaic nymph phenotypes were similar for both ReMOT Control and embryonic microinjection with RNP^w^ in protocol 1. Interestingly, from the 412 G_0_ screened for ReMOT Control, we only found five eye/body color mosaic mutants. We did not observe any live nymph or adult lacking pigmentation in both eyes. The DNA sequences derived from subtle mosaic-eye individuals showed overlapping peaks at the target sites that had to be resolved with ICE analysis. The individuals that showed distinct indels at the target sites either did not survive the embryo or nymphal stages. Moreover, although we did not observe visible mutations in the G_0_ progeny of BAPC-injected females, we were able to infer *indels* in the sequences of G_0_ individuals that showed a wild-type phenotype. Altogether, our data suggest that mutations through embryonic and female injections are not being expressed in the whole organism. This may be due to the timing of injections, the sub-optimized composition of the RNP mix or to operative reasons biasing against the rearing and screening of mutants with better expressivity. Modification to injection mix concentrations, EERs, and other aspects of ACP rearing may improve the chances to generate mutations in the germline during early embryogenesis that can be inherited. Alternatively, following on the previous work with *O. fasciatus,* it is also required to demonstrate if knocking out the *w* gene targeted here confers fitness costs that reduce the chances for survival of such mutants under our rearing conditions (30). Also, it may be possible to use other visible target genes such as orthologs of the *D. melanogaster* genes *scarlet*, *brown* or *vestigial*.

We were not able to demonstrate if the light color variant males observed after the injections with ReMOT Control were off-target mutants. A similar result was observed with *N. vitripennis* for the gene *cinnabar (cin),* in which injections produced eye mutants that did not appear with the typical *cin* phenotype and neither they carried *indels* in the *cin* targeted gene (28).

ReMOT Control uses the peptide ligand “P2C” for ovary transduction of Cas9-RNP. This ligand has been tested to deliver Cas9-RNP in insect species from the orders Hemiptera, Diptera, Coleoptera and Hymenoptera, and in ticks (Arthropoda: Chelicerata) (21,24–28,40). “P2C” is a derived peptide from the Yolk Protein 1 of *D. melanogaster* that has been shown to enhance the entrance of fused proteins into the developing oocytes in the ovaries of mosquitoes (21). There is no homology of P2C to any vitellogenin or vitellogenic protein in *D. citri*, and thus it is surprising that it is functional in this species. Alternatively, our observations suggest that in certain species that do not have the receptors to recognize the P2C ligand, this peptide is not acting as an enhancer for ovary entrance, but instead the RNP is able to enter the oocytes by regular endocytosis. This is supported by previous work using ReMOT Control that demonstrated that Cas9-RNP was able to enter the ovary and produce germline mutagenesis in the oocytes of *Ae aegypt*i and *N.vitripennis* (*21,28*). The same was demonstrated in the mite *Tetranychus urticae* (*22*). Similarly, it was demonstrated recently that the ovaries of *Blatella germanica* and *Tribolium castaneum* also take up Cas9-RNP via endocytosis producing high rates of mutagenesis (26). In the present study, we also demonstrate that our control injections (though limited in number) with Cas9/sgRNA-*w* also produced successful gene editing in *D. citri* with a higher efficiency than using the P2C ligand. Control Cas9 injections were performed without saponin, so it is not possible to directly compare editing efficiency between control injections and ReMOT injections (which included saponin). It is possible that, like in *Bemisia tabaci,* saponin is inhibitory during ReMOT injections, and it would be interesting to evaluate editing efficiency using ReMOT without saponin as efficiency may be significantly improved. However, if the rates of germline gene editing using RNP alone are consistent, this method may be more practical to test other targets and generate heritable mutations that result in one or several mutant *D. citri* lines with different phenotypic markers useful for future biological and genetic studies.

ReMOT Control and BAPC are convenient alternatives for those researchers interested in functional genomics of *D. citri* that lack the expertise or the equipment necessary to carry out embryonic microinjections. However, these methods are yet to be proven to generate integration of DNA into the *D. citri* genome. Therefore it is still vital to continue the optimization of the embryonic microinjection protocols presented here. Nevertheless, we are aware that the ACP presents particular challenges such as the lack of genetic markers and its complexity of rearing which can affect mutagenesis experiments more dramatically compared to other insects for which artificial diet systems have been established. Although these methods have shown that gene editing is plausible, both methods have their setbacks. Embryonic injections are time costly, require specialized equipment, skills, and expertise, and are subject to lower survival rates. On the other hand, female injections are relatively easy to perform and can result in many affected offspring, however, this method requires additional complex crossing procedures and has not yet been applied for transgenesis.

It is also important to progress towards additional steps required to produce and maintain germ line mutants as well as transgenic *D. citri*. First, for both embryo and adult female injection, it is important to recognize the most convenient timing for injections. For instance before cellularization in *D. citri* embryos, or at the peaks of endocytosis in vitellogenic females. Second, the production of an artificial diet as well as an artificial surface for egg lays would make embryo collection much more simple, efficient, and reliable. Finally, alternative approaches for injecting eggs located on the plants to improve the speed and quality of injections would be immensely beneficial to *D. citri* mutagenesis. Potentially using an automated robotic system similar to those reported for *Caenorhabditis elegans*(47) and insect embryos (48) could make efficient injections easier, faster, and more reliable with an easy-to-use software interface.

## Conclusion

In conclusion, we have developed multiple approaches to delivering CRISPR/Cas9 to *D. citri* embryos that have successfully generated somatic mutations. Although this work marks the start of a new era for genetic-based vector control tools for *D. citri*, much work still needs to be done. From our experiments, we found female injections of Cas9 (either ReMOT or just Cas9 RNP) itself was the easiest and most reliable way of generating somatic mutations. However, currently, Cas9 delivery methods based on female injections have not been able to generate transgenic organisms due to the lack of ability to encourage the absorption of donor plasmid into the oocytes. From our work, it is encouraging to confirm that it is possible to generate mutations using embryonic injections. Further investigation into injecting embryos without removing them from the plants could improve survival and thus greater rates of mutagenesis. With this method, the next steps for advancing towards generating genetic-based vector control tools would be to create a heritable mutation, maintain a stable mutant line, and develop transgenesis.

## Methods

### Insect lines and host plants

HLB-free colonies of *D. citri* were a generous gift from the Stouthammer laboratory at the University of California Riverside (UCR). Insects were kept at 26°C and 60% relative humidity in a rearing chamber (Caron^®^ model 6025, Caron, Marietta, O) at the University of California San Diego. We used 25–cm-tall curry plants (*Murraya koenigii*) (gifted from the Stouthammer lab and purchased from Exotica Rare Fruit Nursery (Vista, CA)) and seeds grown into 5-8-cm tall seedlings (DE-Power, Zhongshan, China) gifted from the Stouthammer lab and purchased from David Saim, Long Beach, CA) as an oviposition substrate as well as a feeding source for adults and nymphs. Plants and seedlings were reared in a reflective mylar grow tent (fitted with LED grow lights (Sunraise, Guangdong, China). Plants were watered weekly and fertilized with Osmocote and MiracleGro (ScottsMiracleGro, Marysville, OH) according to manufacturer’s instructions once a month. To perform embryonic microinjections directly on the plants, we used 4–8-week-old seedlings grown individually in drosophila vials (Genesee Scientific, San Diego, CA).

### Target site annotation and sgRNA design and evaluation

To determine the sequences and location of the *white* and *kh* genes in the most recent version of the *D. citri* genome (v3.0), we first ran tBlastn on the website for citrus greening genome resources for *D. citri* (https://citrusgreening.org/tools/blast?db_id=17) using as a query the *D. melanogaster* proteins NP_476787.1 and NP_523651.4 (encoded by the *white* and *kh* genes, NM057439.2 and NM078927.3 respectively). We downloaded the scaffold sequences containing the orthologs and manually located and annotated the exons for each gene using *D. mel* as a reference for the putative mRNA sequences found in NCBI. We used CHOPCHOP to identify the top three sgRNAs with the highest on-target and cleavage efficiency indexes across exons 1–6 of the *white* gene and exons 1 and 2 of the *kh* gene (49). Then, to validate the integrity of the predicted sgRNA target sequences in our insect colonies, we designed primers across both target genes and used them to PCR amplify each target site (see list of primers in **Supplementary Table 1** and the *D. citri* sequences in **Supplementary Figure 2**). The sgRNAs targets that did not show nucleotide variation were tested for cleavage *in vitro* using a plasmid or a PCR product as the template (see below). At the time of the analysis, the *D. citri* genome was not included in any of the platforms for sgRNA design, so we did not run an analysis of potential off-targets.

### SgRNA synthesis

SgRNAs were transcribed *in vitro* and purified using Megascript and MegaClear kits (Thermofisher Scientific, Waltham, MA) following the manufacturer’s instructions. In brief, a PCR template containing the primer CRISPR-R and a CRISPR-F (containing the target sequence and a T7 promoter) were self-annealed and amplified using Phusion^®^ (New England BioLabs, Ipswich, MA) polymerase with the following conditions: 98°C x 2 min, 34 cycles of 98°C x 30 sec, 56°C x 10 sec, 72°C x 10 sec, and a final extension step of 72°C x 5 min. In addition, modified sgRNAs WhEx2.1 and Kh7.1 and Kh7.2 were ordered from Synthego (Redwood City, CA).

### Protein purification and *in vitro* cleavage assay

To deliver the ribonucleoprotein into the oocytes of females, we expressed and purified in-house two different Cas9 derivatives: P2C-mCherry and P2C-Cas9 as previously reported (21,23). We compared the cleavage activity of P2C-Cas9 with commercially available Cas9 from PNABio (Thousand Oaks, CA).

### Embryonic microinjection protocols

To collect *D. citri* eggs for embryonic microinjections, we prepared an oviposition cage with one or two curry plants with abundant flushes and introduced 20–30 gravid ACP females aged 7–10 days. Plants were monitored every two hours for eggs. When eggs were found, the plant was cleared of adult *D. citri* and removed from the cage. Eggs were then collected by gently removing them from the curry leaves with a teasing needle. We then prepared a surface for alignment using filter paper moistened with ultrapure water mounted on a glass slide. A second filter paper was overlaid onto the first to create a vertical wall along which the eggs were aligned against. We prepared quartz needles on a P2000 needle puller using the protocol: Heat: 650, Fil: 4, Vel: 40, Del: 150, Pul: 170. To optimize embryonic microinjections in *D. citri*, we developed two different protocols. Protocol 1, wherein we injected embryos removed from the plants, and Protocol 2, wherein we injected embryos on the plants (**Fig. 2**). Females aged 7–10 days were allowed to lay eggs on large/older plants (protocol 1) or seedlings (protocol 2). For protocol 1, eggs were removed from the large plants with a dissecting needle and transferred to a paper filter wetted with UltraPure distilled water (**Fig. 2A**). Eggs were aligned to facilitate injection at the posterior pole, the wider region with a distinctive dark yellow color that is opposite to the narrow tip of the egg (**Fig. 2B**). For injections, we used freshly pulled quartz needles and a FemtoJet 4i (Eppendorf, Hamburg, Germany) at Pc value of 100. Because the needles often clog, we typically prepare 10 needles for each session. Following injections, the injected eggs were transferred to agar plates to humidify the developing embryos (**Fig. 2C**). Once the nymphs hatched, nymphs were transferred to large plants for development.

For protocol 2, we allowed the females to lay between 10-20 eggs on young seedlings (2-3 months old) that are planted in vials containing a mixture of vermiculite and perlite. The seedlings are easily removed from the soil and mounted onto a microscope slide using tape (**Fig 2D**). The locations of the eggs are identified under a dissecting scope and tape was used to keep the area accessible to the needles. Eggs were injected at the nearest point to the posterior pole (**Fig 2E**). Eggs not accessible to injections were removed from the plant or killed. After injections, seedlings were transferred to a growth chamber to allow for the growth of both the plants and nymphs that hatch on the plant (**Figure 2F**).

### Female microinjection protocols

Curry plants containing 4th and 5th instar nymphs were placed into a large cage. The plants were inspected and cleared of all adults. Every following day, newly emerged adults were collected and placed into enclosed cups with seedlings (**Figure S3B**). The adults were then aged until 7-10 days post eclosion. To prepare for injections, aged adults were anesthetized using CO2 or ice. Anesthetized adults were sex separated. Males were placed onto collection tube lids with moistened paper towels. They were then placed back into the original seedling cup. Females were injected with the injection mixture using a pulled microcapillary needle connected to an aspirator tube. Needles were inserted in a membranous region between sternites two and three. Injected females were then placed back into the cup in the same manner as the males. Following injections, collection tube lids were observed for the number of females that did not survive the injection procedure. Every day following, the plants were checked for eggs and nymphs. Five days post-injection, injected adults were collected, counted, and placed into a new cage with fresh flushing plants while the previous plants with eggs were transferred to a new cage. Eggs were allowed to develop and emerge. Once nymphs were 4th to 5th instar stage, they were removed using an embryo pick and recorded for any mutant phenotypes. Identified mutants were outcrossed and allowed to produce G1 progeny. All nymphs and adults were placed into PCR tubes with squish buffer for DNA extraction and sequencing.

### Determination and Confirmation of Mutants

Emerged nymphs were collected and observed under a stereoscope for potential mutant phenotypes. Each nymphs was collected and stored in their own 0.2mL PCR tube and labeled as a potential mutant or wild-type. Genomic DNA was then extracted from each nymph. The relevant target sequence was then PCR amplified. The PCR amplified product was run through a 4% agarose gel to visually inspect for amplified products of different sizes. The bands were excised and DNA was extracted from them and sent for sanger sequencing. Sequences were observed for INDELs or perturbations around the target site. Further ICE analysis was performed to confirm potential INDELs. Nymphs identified as potential mutants were confirmed to have PCR products that resulted in multiple bands which were confirmed to contain INDELs through sanger sequencing and ICE analysis.

## Statistical analysis

In all experiments, a minimum of three replicates were used to make comparisons between means. Unpaired t-tests were performed using GraphPad Prism version 8.0.0 for Windows, GraphPad Software, San Diego, California USA, www.graphpad.com.

## Acknowledgments

We thank Sen Miao, Steven Olivas, and Richard Stouthammer for their expertise on *D. citri* husbandry as well as their contribution of *D. citri and M. koenigii* seeds and seedlings. Figures were created with www.BioRender.com.

## Author Contributions

O.S.A., conceived and designed the experiments. D.C.R., C.L.G., and M.B., performed molecular and genetic experiments. J.L.R. provided ReMOT reagents. All authors contributed to the writing, analyzed the data, and approved the final manuscript.

## Ethical conduct of research

All animals were handled in accordance with the Guide for the Care and Use of Laboratory Animals as recommended by the National Institutes of Health and approved by the UCSD Biological Use Authorization (BUA #R2401).

## Supplementary Information

1. *white* gene genomic DNA sequence encompassing the sgRNA target sites and its corresponding aminoacid sequence from laboratory colonies of *D. citri*.

A. DNA Sequence for the *w* gene from our laboratory colonies used to design sgRNA. Exons are marked in colors starting with exon 2 and finishing with exon 9. sgRNA target sites are marked with red font.

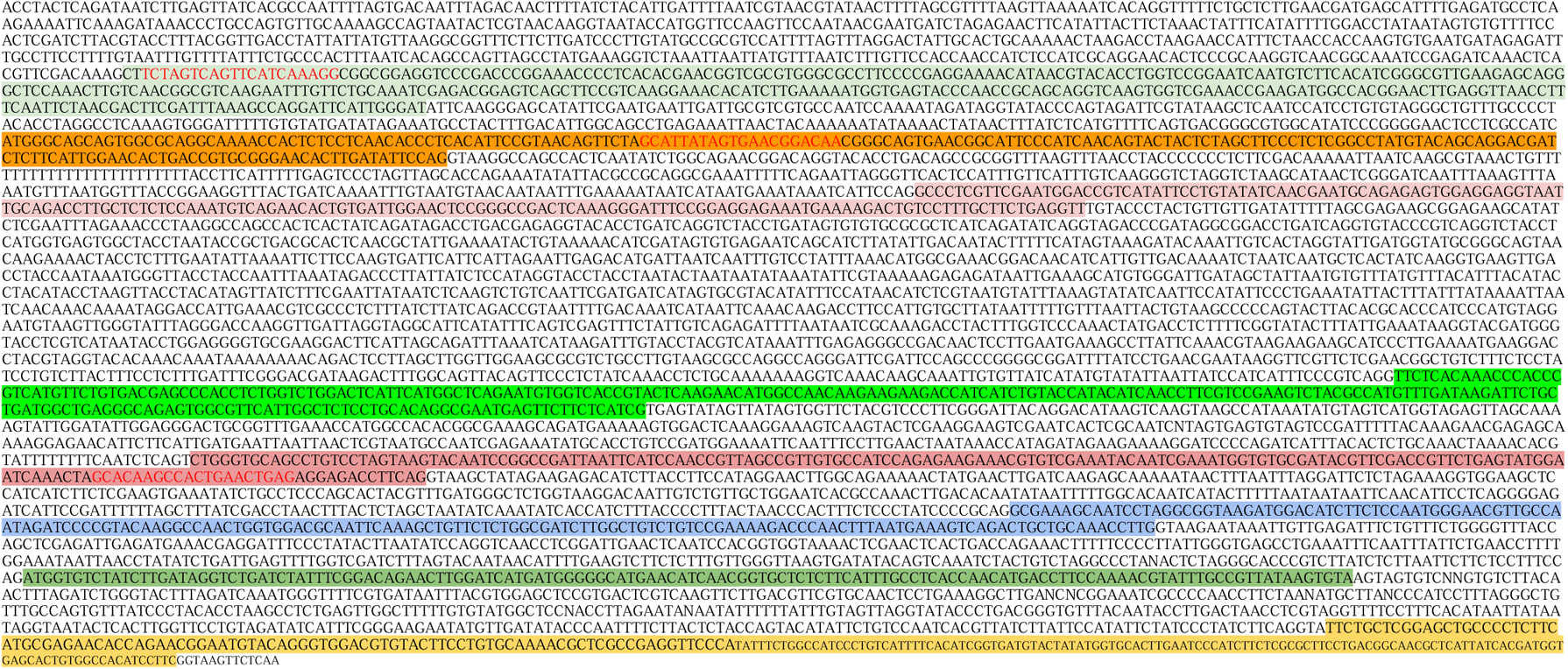

B. Corresponding aminoacid sequence for the *w* gene from *D. citri* lab colonies. sgRNA target sites 2.1, 3.1 and 6.1 marked with color font.

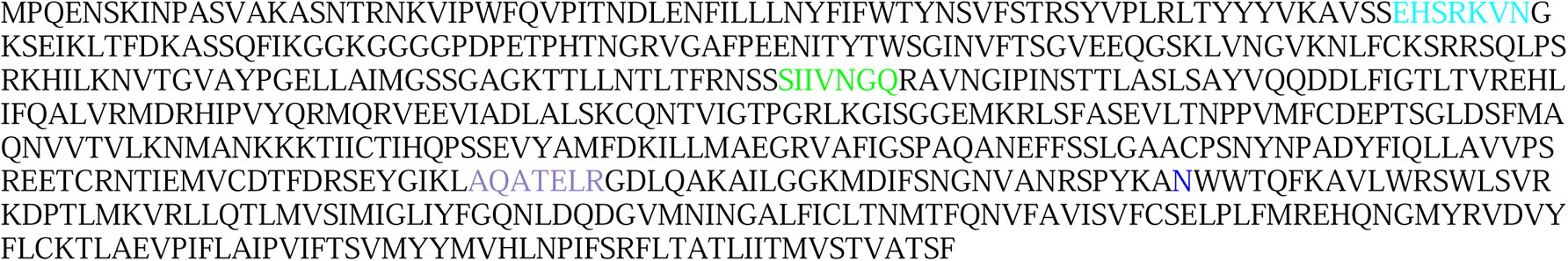

2. *kh* gene genomic DNA sequence encompassing the sgRNA target sites and its corresponding aminoacid sequence from laboratory colonies of *D. citri*.

A. DNA Sequence for the *kh* gene from our laboratory colonies to design sgRNA. Exons are marked in colors starting with exon 7 and finishing with exon 10. sgRNA target sites are marked with red font.

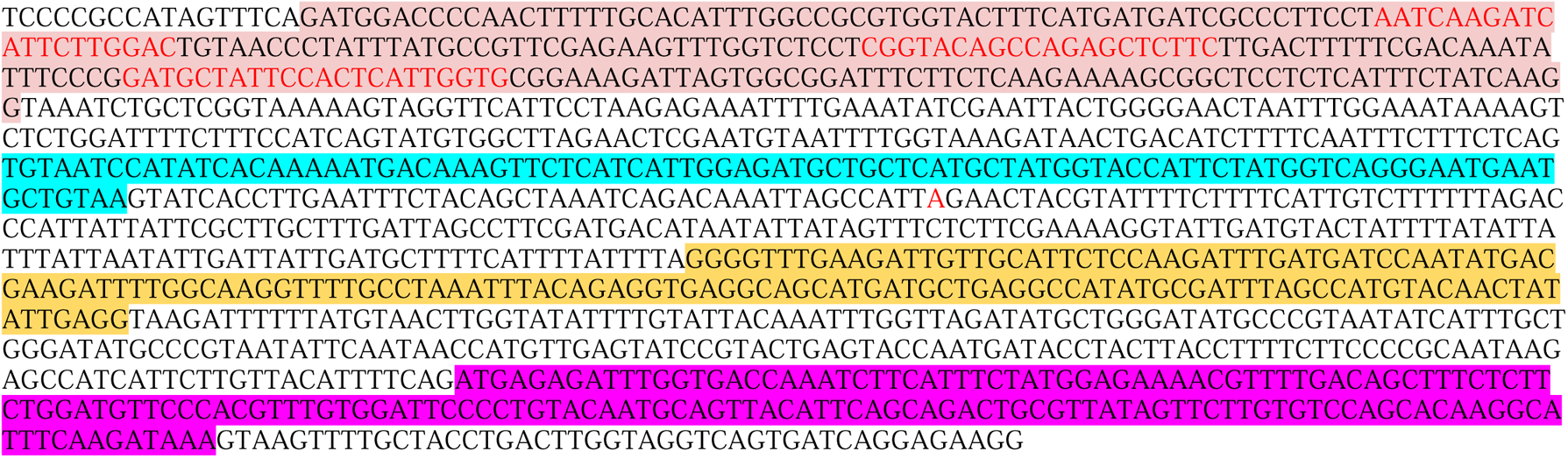

B. Corresponding aminoacid sequence for the *w* gene from *D. citri* lab colonies. sgRNA target sites 7.1, 7.2 and 7.3 are marked with color font.

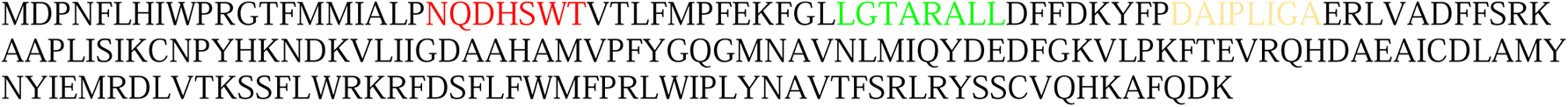

**Supplementary Protocol 1. Embryonic injections of *D. citri* eggs removed from curry plants.**

After allowing mature females to lay eggs within the flush of a curry plant, a dissecting teasing needle, with a curved bend at the tip, was used to gently lift and pluck the eggs from the plant. The removed eggs were placed and collected on 5.6% gelatin plates. Once collected, the eggs were arranged on top of the bottom square of two squares of overlapping moistened filter paper both placed on top of a glass microscope slide. The eggs were aligned against the edge of the upper filter paper square so the more rounded posterior end of the egg was accessible for injections (Fig. 2A). The aligned eggs were then injected with Cas9/sgRNA RNP using pulled quartz capillary needles on a P-2000 needle puller, pulled using the following parameters program 77 (Heat: 650, Fil: 4, Vel: 40, Del: 150, Pull: 170), and a FemtoJet 4i set to a Pi value of 600–1000 and a Pc value of 150–300, depending on the opening size of the needle. As injections continue, the needle tip may break or clog. To accommodate, the pressure was adjusted to maintain the volume injected. To address full needle clogging, a major issue with many insect embryo injections, we prepared 10 needles before each session, loading them with 3–4 ul of the mix when needed. To prevent desiccation of the eggs from the heat from the light sources, Ultrapure water was carefully added to the filter paper with a bulb pipette as needed. Once all eggs were injected, the filter paper with the injected eggs was transferred to fresh gelatin plates. Eggs were transferred to fresh gelatin plates every two days to reduce mold growth.

**Supplementary Protocol 2. Embryonic injections of *D. citri* eggs directly on curry plants**

Young *Murraya koenigii* seedlings were grown from seeds in plastic vials for 4-8 weeks until they were 2-3 inches in height. Seedlings presenting suitable foliage for *D. citri* oviposition were presented to mature females. Seedlings were observed every 30 min for eggs. Once containing eggs, seedlings were uprooted and mounted onto a microscope slide using tape (Fig. 2). The locations of the eggs were determined using a dissecting scope, and foliage that interfered with the needle path was either taped down or carefully removed. Eggs were injected as close to the posterior pole as feasible (Fig. 2). Eggs that were not accessible for injection were removed from the plant or destroyed. After injection, seedlings were transferred to an ACP growth chamber to allow the eggs to develop and eclose (Fig. 2).

**Supplementary Protocol 3. Adult Female Injection**

For adult female injections, insects aged 7–10 days post eclosion were collected and anesthetized using either ice or CO_2_. Once anesthetized insects were placed on either a portable chill table (1429, BioQuip, Rancho Dominguez, CA) or a CO_2_ pad (59-172, Genesee Scientific, San Diego, CA) and sorted by sex. Males were collected onto dishes lined with filter paper moistened with ddH_2_O and placed back into the cage. Females were kept on either the chill table or CO2 pad during injections. A quartz capillary needle filled with 10 µL of injection mixture was attached to an aspirator tube (A5177-5EA, Sigma Aldrich, Munich, Germany). Females were injected until slightly inflated into the lateral region between their abdomen and thorax, which is less sclerotized compared to the rest of the *D. citri* body for easier penetration with reduced needle breakage. Injected females were gently moved to a dish lined with filter paper moistened with ddH2O. The collection dish was then placed into a cage containing fresh curry plants and the males previously collected and separated.

## Supplementary Figures

**Figure S1.**
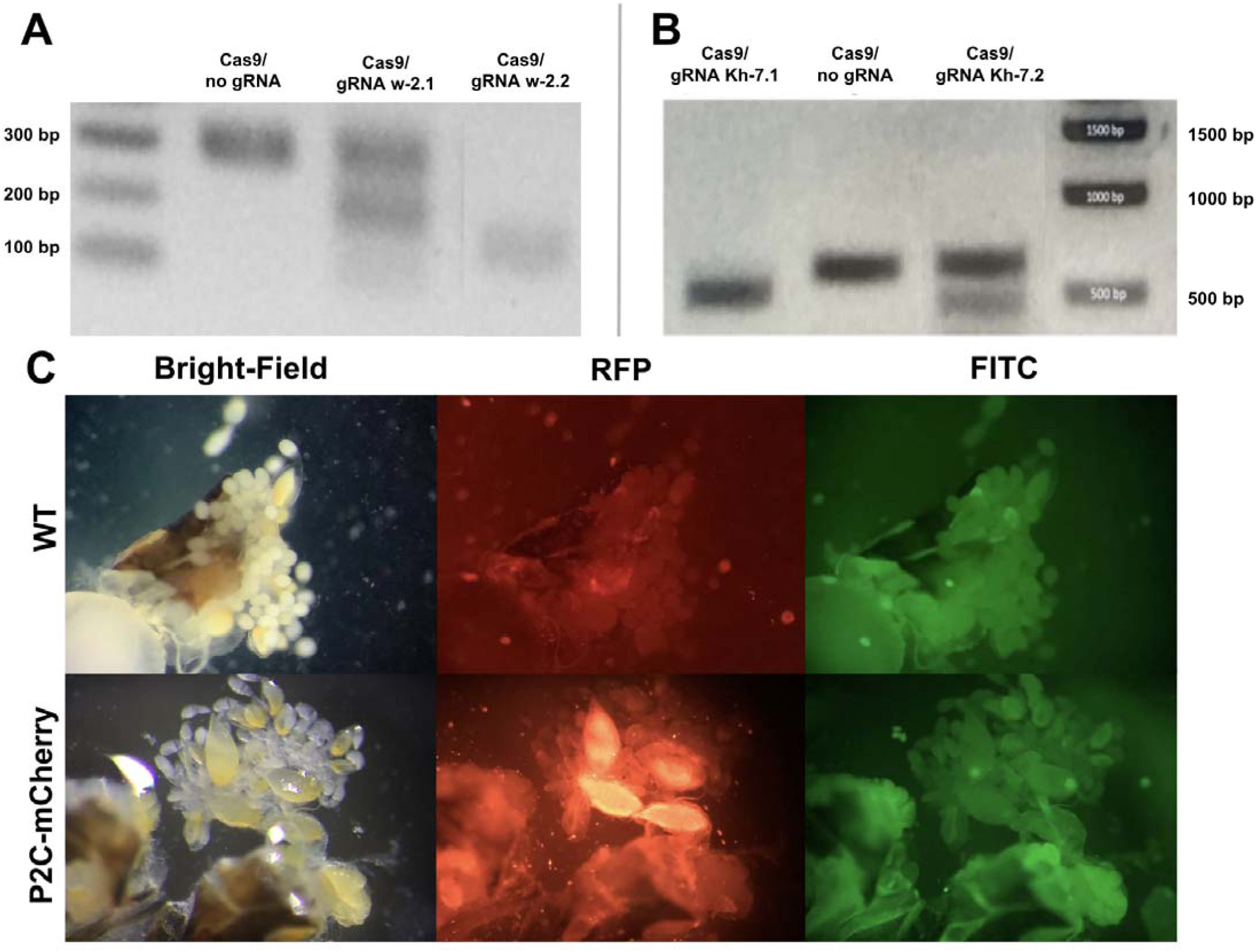
Testing cleavage ability of sgRNAs targeting *w* and *kh* and localization of P2C protein. **A.** *In vitro* cleavage assay of *D. citri* PCR products with gRNAs *w*-2.1 and *w*-2.2. **B.** *In vitro* cleavage assay of *D. citri* PCR products with gRNAs *kh*-7.1 and *kh*-7.2. **C.** Ovaries and embryos of *D.citri* female injected with the protein P2C-mCherry and imaged under Bright-field, RFP and FITC filters.

**Figure S2.**
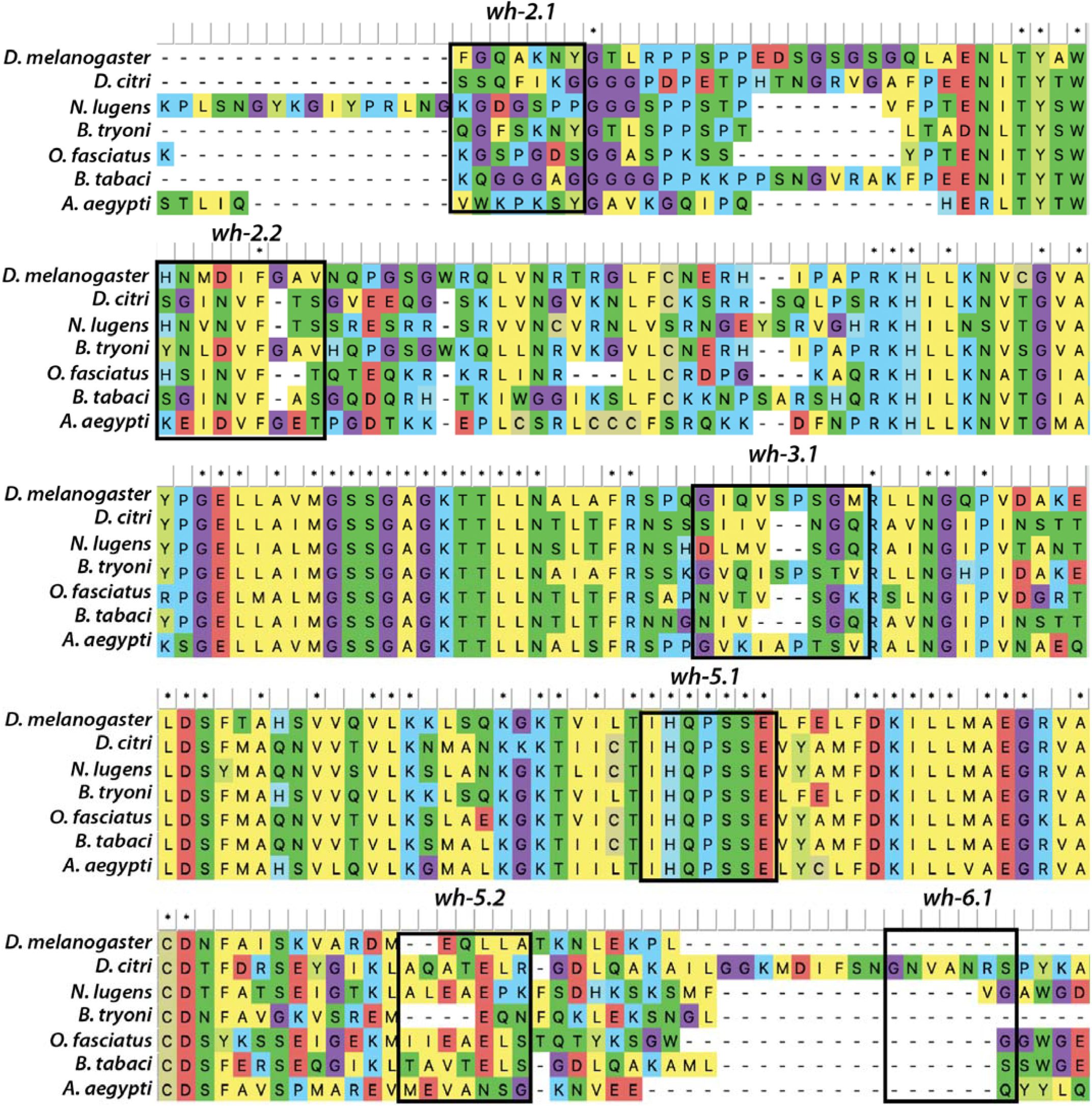
Protein sequence homology of *w* between *D. citri*, *D. mel* and other hemipterans with known *w* mutants. Homology of the *w* gene across the species, *Drosophila melanogaster, Diaphorina citri, Nilaparvata lugens, Bactrocera tryoni, Oncopeltus fasciatus, Bernisia tabaci, and Aedes aegypti*. Amino acid sequences pertaining to designed sgRNAs are outlined in boxes and labeled accordingly.

**Figure S3.**
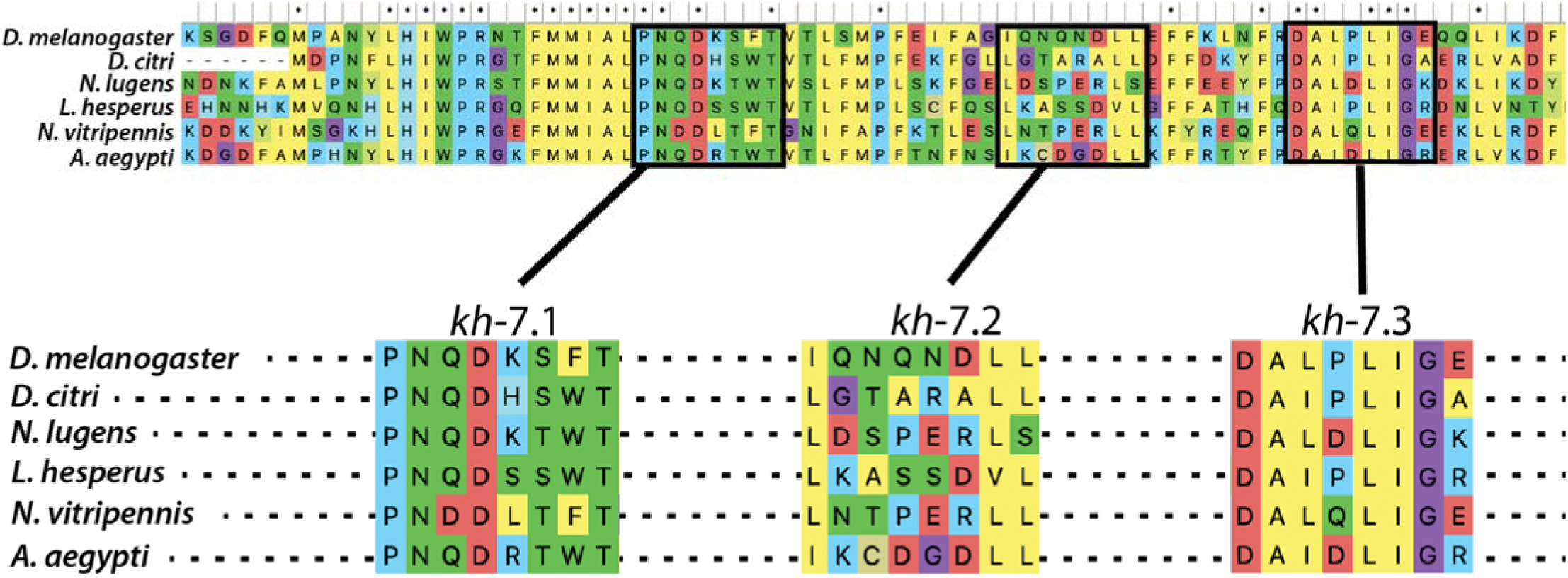
Protein sequence homology of *kh* between *D. citri*, *D. mel* and other hemipterans with known *kh* mutants. A. Homology of the *kh* gene across the species: *D. melanogaster, D. citri, Ae aegypti, N. vitripennis, N. lugens, and L. hesperus.*Amino acid sequences pertaining to designed sgRNAs are outlined in boxes and labeled accordingly.

**Figure S3.**
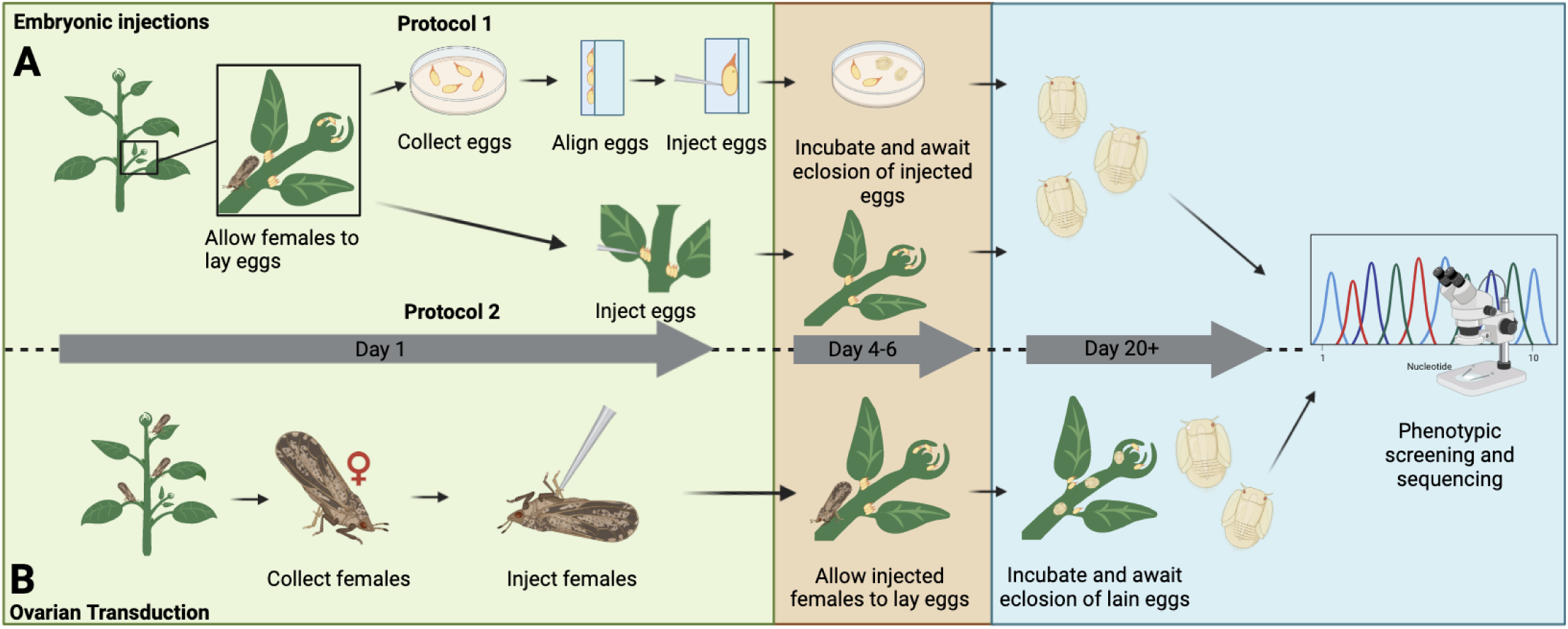
Embryonic and adult female injection protocols. A) Embryonic injections were performed by either carefully removing eggs from the plant and aligning them onto a glass slide to be injected on the posterior end or injecting the eggs while still on the plant. B) For adult female injections, adult gravid females were collected, anesthetized using CO_2,_ and injected between their thorax and abdomen using a microneedle attached to a mouth pipette. Females were then allowed to recover and were introduced to plants for egg-laying. Once the eggs hatched, the resulting nymphs were screened and sequenced for the expected phenotypes. Created with BioRender.com.

**Figure S4.**
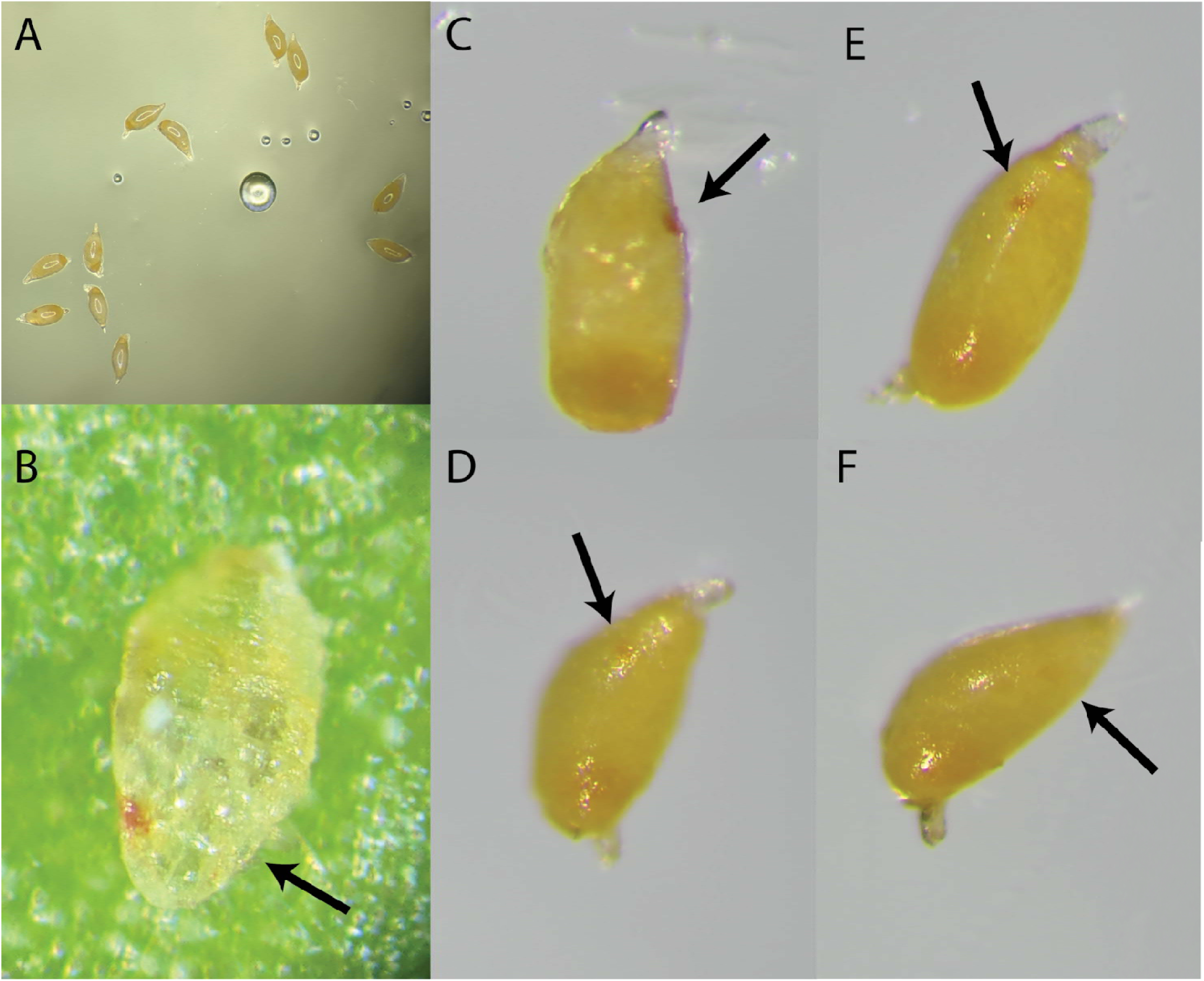
Phenotypes of developing embryos after injection with RNP^w^. **A.** Post-injection embryos resting and developing on artificial media plates. **B.** 1st instar nymph with one mutant eye phenotype. The nymph desiccated and died post transferring from artificial media plate to young curry flush leaf. **C-F.** Developing embryos with varied eyespot color intensities. All embryos were laid and injected at the same time.

**Figure S5.**
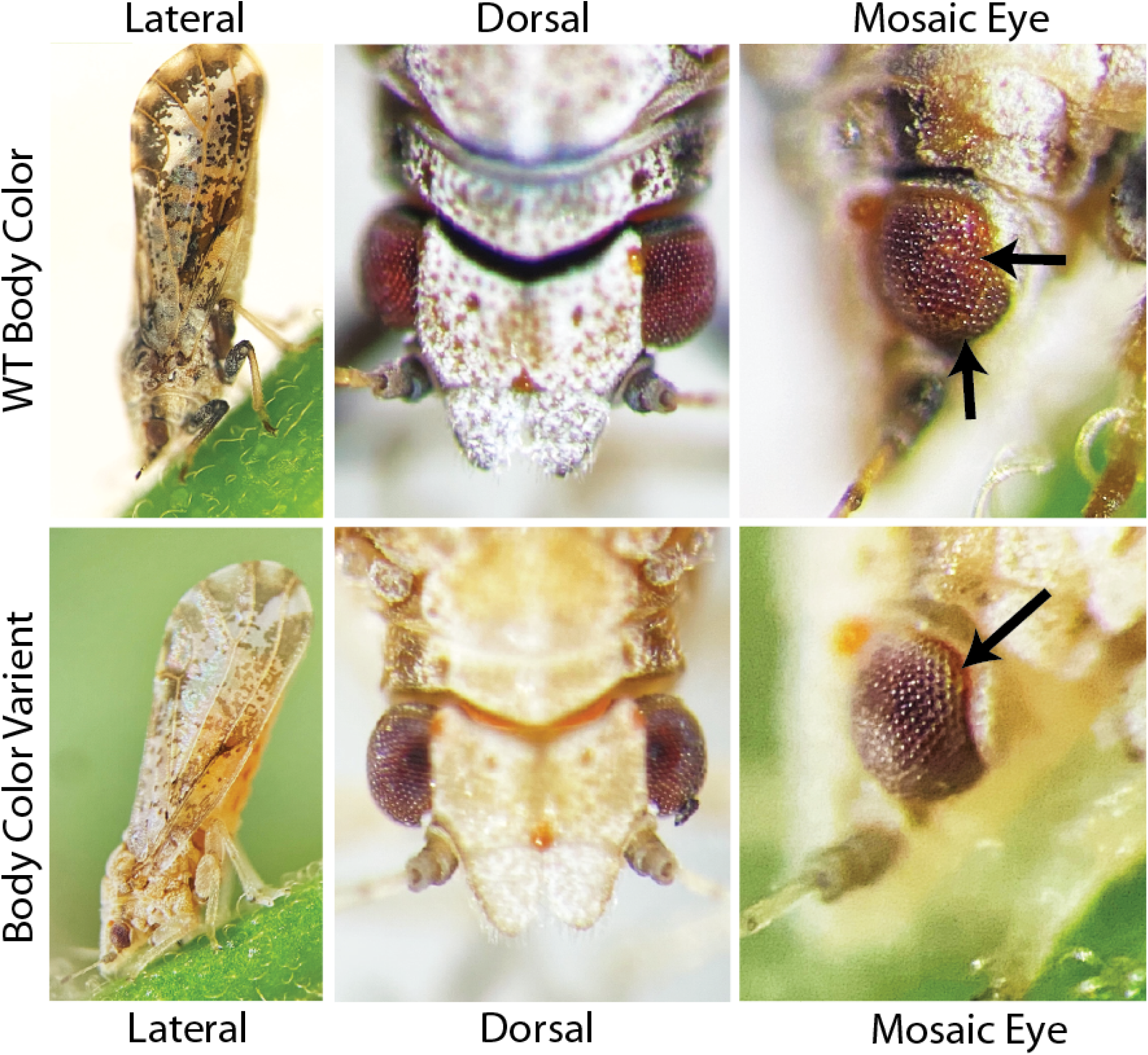
Body color variability of G_0_ adults obtained from ReMOT Control. Comparison of body color of a mosaic eye male (top) and male with light body color obtained after injection with ReMOT Control. The variant individual did not show indels at the target site but was not observed in our colonies or experimental treatments.

## References

1. Hall DG, Richardson ML, Ammar ED, Halbert SE. Asian citrus psyllid, Diaphorina citri, vector of citrus huanglongbing disease. Entomol Exp Appl. 2013 Feb 7;146(2):207–23.

2. Bassanezi RB, Montesino LH, Stuchi ES. Effects of huanglongbing on fruit quality of sweet orange cultivars in Brazil [Internet]. Vol. 125, European Journal of Plant Pathology. 2009. p. 565–72. Available from: http://dx.doi.org/10.1007/s10658-009-9506-3

3. Graham J, Gottwald T, Setamou M. Status of Huanglongbing (HLB) outbreaks in Florida, California and Texas [Internet]. Vol. 45, Tropical Plant Pathology. 2020. p. 265–78. Available from: http://dx.doi.org/10.1007/s40858-020-00335-y

4. Lee JA, Halbert SE, Dawson WO, Robertson CJ, Keesling JE, Singer BH. Asymptomatic spread of huanglongbing and implications for disease control. Proc Natl Acad Sci U S A. 2015 Jun 16;112(24):7605–10.

5. Gottwald TR. Current epidemiological understanding of citrus Huanglongbing. Annu Rev Phytopathol. 2010;48:119–39.

6. Polek M, Vidalakis G, Godfrey K. Citrus Bacterial Canker Disease and Huanglongbing (Citrus Greening) [Internet]. 2007. Available from: http://dx.doi.org/10.3733/ucanr.8218

7. Bové JM. HUANGLONGBING: A DESTRUCTIVE, NEWLY-EMERGING, CENTURY-OLD DISEASE OF CITRUS. J Plant Pathol. 2006;88(1):7–37.

8. Munir S, He P, Wu Y, He P, Khan S, Huang M, et al. Huanglongbing Control: Perhaps the End of the Beginning. Microb Ecol. 2018 Jul;76(1):192–204.

9. Li X, Ruan H, Zhou C, Meng X, Chen W. Controlling Citrus Huanglongbing: Green Sustainable Development Route Is the Future. Front Plant Sci. 2021 Nov 15;12:760481.

10. Alquézar B, Carmona L, Bennici S, Miranda MP, Bassanezi RB, Peña L. Cultural Management of Huanglongbing: Current Status and Ongoing Research. Phytopathology. 2022 Jan;112(1):11–25.

11. Tiwari S, Killiny N, Stelinski LL. Dynamic insecticide susceptibility changes in Florida populations of Diaphorina citri (Hemiptera: Psyllidae). J Econ Entomol. 2013 Feb;106(1):393–9.

12. Tiwari S, Mann RS, Rogers ME, Stelinski LL. Insecticide resistance in field populations of Asian citrus psyllid in Florida. Pest Manag Sci. 2011 Oct;67(10):1258–68.

13. Parasitism of Diaphorina citri (Hemiptera: Liviidae) by Tamarixia radiata (Hymenoptera: Eulophidae) on residential citrus in Texas: Importance of colony size and instar composition. Biol Control. 2022 Feb 1;165:104796.

14. Density dependent mortality, climate, and Argentine ants affect population dynamics of an invasive citrus pest, Diaphorina citri, and its specialist parasitoid, Tamarixia radiata, in Southern California, USA. Biol Control. 2021 Aug 1;159:104627.

15. Yang Y, Huang M, Beattie GAC, Xia Y, Ouyang G, Xiong J. Distribution, biology, ecology and control of the psyllid Diaphorina citri Kuwayama, a major pest of citrus: A status report for China. Int J Pest Manage [Internet]. 2007 Feb 23 [cited 2022 Sep 16]; Available from: https://www.tandfonline.com/doi/abs/10.1080/09670870600872994

16. Hammond A, Galizi R, Kyrou K, Simoni A, Siniscalchi C, Katsanos D, et al. A CRISPR-Cas9 gene drive system targeting female reproduction in the malaria mosquito vector Anopheles gambiae. Nat Biotechnol. 2016 Jan;34(1):78–83.

17. Buchman A, Gamez S, Li M, Antoshechkin I, Li HH, Wang HW, et al. Broad dengue neutralization in mosquitoes expressing an engineered antibody. PLoS Pathog. 2020 Jan;16(1):e1008103.

18. Kandul NP, Liu J, Sanchez C HM, Wu SL, Marshall JM, Akbari OS. Transforming insect population control with precision guided sterile males with demonstration in flies. Nat Commun. 2019 Jan 8;10(1):84.

19. Burckhardt D. Psylloid pests of temperate and subtropical crop and ornamental plants (Hemiptera, Psylloidea): a review. Trends in Agricultural Sciences, Entomology. 1994;2:173–86.

20. Hunter WB, Gonzalez MT, Tomich J. BAPC-assisted CRISPR/Cas9 System: Targeted Delivery into Adult Ovaries for Heritable Germline Gene Editing (Arthropoda: Hemiptera) [Internet]. bioRxiv. 2018 [cited 2020 Jul 29]. p. 478743. Available from: https://www.biorxiv.org/content/10.1101/478743v1.abstract

21. Chaverra-Rodriguez D, Macias VM, Hughes GL, Pujhari S, Suzuki Y, Peterson DR, et al. Targeted delivery of CRISPR-Cas9 ribonucleoprotein into arthropod ovaries for heritable germline gene editing. Nat Commun. 2018 Aug 1;9(1):3008.

22. Dermauw W, Jonckheere W, Riga M, Livadaras I, Vontas J, Van Leeuwen T. Targeted mutagenesis using CRISPR-Cas9 in the chelicerate herbivore Tetranychus urticae. Insect Biochem Mol Biol. 2020 May;120:103347.

23. Heu CC, McCullough FM, Luan J, Rasgon JL. CRISPR-Cas9-Based Genome Editing in the Silverleaf Whitefly (). CRISPR J. 2020 Apr;3(2):89–96.

24. Macias VM, McKeand S, Chaverra-Rodriguez D, Hughes GL, Fazekas A, Pujhari S, et al. Cas9-Mediated Gene-Editing in the Malaria Mosquito by ReMOT Control. G3. 2020 Apr 9;10(4):1353–60.

25. Shirai Y, Daimon T. Mutations in cardinal are responsible for the red-1 and peach eye color mutants of the red flour beetle Tribolium castaneum. Biochem Biophys Res Commun. 2020 Aug 20;529(2):372–8.

26. Shirai Y, Piulachs MD, Belles X, Daimon T. DIPA-CRISPR is a simple and accessible method for insect gene editing. Cell Rep Methods. 2022 May 23;2(5):100215.

27. Li X, Xu Y, Zhang H, Yin H, Zhou D, Sun Y, et al. ReMOT Control Delivery of CRISPR-Cas9 Ribonucleoprotein Complex to Induce Germline Mutagenesis in the Disease Vector Mosquitoes Culex pipiens pallens (Diptera: Culicidae). J Med Entomol. 2021 May 15;58(3):1202–9.

28. Chaverra-Rodriguez D, Dalla Benetta E, Heu CC, Rasgon JL, Ferree PM, Akbari OS. Germline mutagenesis of Nasonia vitripennis through ovarian delivery of CRISPR-Cas9 ribonucleoprotein. Insect Mol Biol. 2020 Dec;29(6):569–77.

29. Gerard Terradas, Vanessa M. Macias, Hillary Peterson, Sage McKeand, Grzegorz Krawczyk, Jason L. Rasgon. Receptor-Mediated Ovary Transduction of Cargo – ReMOT Control: a Comprehensive Review and Detailed Protocol for Implementation. Transgenic Insects: Techniques and Applications. 2022 Nov 3;125–48.

30. Reding K, Pick L. High-Efficiency CRISPR/Cas9 Mutagenesis of the *white* Gene in the Milkweed Bug *Oncopeltus fasciatu*s [Internet]. Vol. 215, Genetics. 2020. p. 1027–37. Available from: http://dx.doi.org/10.1534/genetics.120.303269

31. Pacheco I de S, de Souza Pacheco I, Doss ALA, Vindiola BG, Brown DJ, Ettinger CL, et al. Efficient CRISPR/Cas9-mediated genome modification of the glassy-winged sharpshooter Homalodisca vitripennis (Germar) [Internet]. Vol. 12, Scientific Reports. 2022. Available from: http://dx.doi.org/10.1038/s41598-022-09990-4

32. Choo A, Crisp P, Saint R, O’Keefe LV, Baxter SW. CRISPR /Cas9LJmediated mutagenesis of the *white* gene in the tephritid pest *Bactrocera tryoni* [Internet]. Vol. 142, Journal of Applied Entomology. 2018. p. 52–8. Available from: http://dx.doi.org/10.1111/jen.12411

33. Heu CC, Gross RJ, Le KP, LeRoy DM, Fan B, Hull JJ, et al. CRISPR-mediated knockout of cardinal and cinnabar eye pigmentation genes in the western tarnished plant bug. Sci Rep. 2022 Mar 22;12(1):4917.

34. Jasinskiene N, Juhn J, James AA. Microinjection of A. aegypti embryos to obtain transgenic mosquitoes. J Vis Exp. 2007 Jul 4;(5):219.

35. Li M, Bui M, Akbari OS. Embryo Microinjection and Transplantation Technique for Nasonia vitripennis Genome Manipulation. J Vis Exp [Internet]. 2017 Dec 25;(130). Available from: http://dx.doi.org/10.3791/56990

36. Santos-Ortega Y, Killiny N. In vitro egg hatching of Diaphorina citri, the vector of Huanglongbing. Entomol Exp Appl. 2020 Nov;168(11):851–6.

37. Xue WH, Xu N, Yuan XB, Chen HH, Zhang JL, Fu SJ, et al. CRISPR/Cas9-mediated knockout of two eye pigmentation genes in the brown planthopper, Nilaparvata lugens (Hemiptera: Delphacidae). Insect Biochem Mol Biol. 2018 Feb;93:19–26.

38. Kotwica-Rolinska J, Chodakova L, Chvalova D, Kristofova L, Fenclova I, Provaznik J, et al. CRISPR/Cas9 Genome Editing Introduction and Optimization in the Non-model Insect Pyrrhocoris apterus. Front Physiol. 2019 Jul 15;10:891.

39. Mankin RW, Rohde B. Mating behavior of the Asian citrus psyllid. In: Asian citrus psyllid: biology, ecology and management of the Huanglongbing vector. UK: CABI; 2020. p. 30–42.

40. Sharma A, Pham MN, Reyes JB, Chana R, Yim WC, Heu CC, et al. Cas9-Mediated Gene-Editing in the Black-Legged Tick, Ixodes Scapularis, by Embryo Injection and ReMOT Control [Internet]. 2020 [cited 2021 Feb 16]. Available from: https://papers.ssrn.com/abstract=3691041

41. Mathew D, Prasad MC. Multiple shoot and plant regeneration from immature leaflets of in vitro origin in curryleaf (Murraya koenigii Spreng). Indian Journal of Plant Physiology. 2007;12(1):18.

42. Paul S, Dam A, Bhattacharyya A, Bandyopadhyay TK. An efficient regeneration system via direct and indirect somatic embryogenesis for the medicinal tree Murraya koenigii. Plant Cell Tissue Organ Cult. 2011 May 1;105(2):271–83.

43. Bhuyan AK, Pattnaik S, Chand PK. Micropropagation of Curry Leaf Tree [Murraya koenigii (L.) Spreng.] by axillary proliferation using intact seedlings. Plant Cell Rep. 1997 Sep;16(11):779–82.

44. Sakthivel A, Velmurugan S, Selvi S, Senthil A. Studies on vegetative propagation in curry leaf (Murraya koenigii Spreng.) [Internet]. [cited 2023 Apr 29]. Available from: https://www.thepharmajournal.com/archives/2021/vol10issue10/PartAK/10-10-271-721.pdf

45. Fuchs H, Bachran C, Flavell DJ. Diving through Membranes: Molecular Cunning to Enforce the Endosomal Escape of Antibody-Targeted Anti-Tumor Toxins. Antibodies. 2013 Apr 17;2(2):209–35.

46. Dalla Benetta E, Chaverra-Rodriguez D, Rasgon JL, Akbari OS. Pupal and Adult Injections for RNAi and CRISPR Gene Editing in Nasonia vitripennis. J Vis Exp [Internet]. 2020 Dec 4;(166). Available from: http://dx.doi.org/10.3791/61892

47. Gilleland CL, Falls AT, Noraky J, Heiman MG, Yanik MF. Computer-Assisted Transgenesis of Caenorhabditis elegans for Deep Phenotyping. Genetics. 2015 Sep;201(1):39–46.

48. Gilleland CL. System and method for high-throughput precision robotic embryo manipulation under computer control at large-scale with high reliability microdosing. 2022 May 27; PCT Patent WO2022251664A2. Available from: https://patents.google.com/patent/WO2022251664A2/en

49. Labun K, Montague TG, Krause M, Torres Cleuren YN, Tjeldnes H, Valen E. CHOPCHOP v3: expanding the CRISPR web toolbox beyond genome editing. Nucleic Acids Res. 2019 Jul 2;47(W1):W171–4.

